# Modeling the demography of species providing extended parental care: A capture-recapture multievent model with a case study on Polar Bears (*Ursus maritimus*)

**DOI:** 10.1101/596437

**Authors:** Sarah Cubaynes, Jon Aars, Nigel G. Yoccoz, Roger Pradel, Øystein Wiig, Rolf A Ims, Olivier Gimenez

## Abstract

1. In species providing extended parental care, one or both parents care for altricial young over a period including more than one breeding season. We expect large parental investment and long-term dependency within family units to cause high variability in life trajectories among individuals with complex consequences at the population level. So far, models for estimating demographic parameters in free-ranging animal populations mostly ignore extended parental care, thereby limiting our understanding of its consequences on parents and offspring life histories.
2. We designed a capture-recapture multi-event model for studying the demography of species providing extended parental care. It handles statistical multiple-year dependency among individual demographic parameters grouped within family units, variable litter size, and uncertainty on the timing at offspring independence. It allows to evaluate trade-offs among demographic parameters, the influence of past reproductive history on the caring parent survival status, breeding probability and litter size probability, while accounting for imperfect detection of family units. We assess the model performances using simulated data, and illustrate its use with a long-term dataset collected on the Svalbard polar bears (*Ursus maritimus*).
3. Our model performed well in terms of bias and mean square error and in estimating demographic parameters in all simulated scenarios, both when offspring departure probability from the family unit occurred at a constant rate or varied during the field season depending on the date of capture. For the polar bear case study, we provide estimates of adult and dependent offspring survival rates, breeding probability and litter size probability. Results showed that the outcome of the previous reproduction influenced breeding probability.
4. Overall, our results show the importance of accounting for i) the multiple-year statistical dependency within family units, ii) uncertainty on the timing at offspring independence, and iii) past reproductive history of the caring parent. If ignored, estimates obtained for breeding probability, litter size, and survival can be biased. This is of interest in terms of conservation because species providing extended parental care are often long-living mammals vulnerable or threatened with extinction.

## INTRODUCTION

Parental care includes any pre-natal and post-natal allocation, such as feeding and protecting the young, which benefits the offspring development and survival chances, thereby enhancing the parent’s reproductive success (Trivers 1972). Altricial mammals having offspring that need to learn complex skills to ensure survival beyond independence, such as hunting, orientation, or nest building, show extended parental care (hereafter EPC; Clutton-Brock 1991). It is defined as a prolonged period, i.e. lasting more than one breeding season, over which one or both parents care for one or several dependent young. This period typically lasts for several years and can extend until lifelong maternal care in primates (Van Noordwijk 2012). For the offspring, the quality and quantity of care received can have long-lasting effects on future survival (e.g. Pavard and Branger 2012), social status (e.g. Shenk and Scelza 2012) and reproduction (Royle et al. 2012). For the parent, investment in one offspring can compromise its own condition or survival and/or its ability to invest in other offspring (siblings or future offspring) (Williams 1966, Stearns 1992). It can indeed take several years during which a parent caring for its offspring will not be available to reproduce, sometimes not until the offspring have reached independence, e.g. on average 2.5 years for female polar bears (Ramsay and Stirling 1988), 3.5 to 6 years for female African elephants (Lee and Moss 1986), and 9.3 years for female Sumatran orangutans (Wich et al. 2004). The fitness costs of losing one offspring, in terms of lost investment and skipped breeding opportunities, are particularly high if death occurs near independence. We therefore expect EPC, through large parental investment and multiple-year dependency among individuals within family units, to cause high variability in life trajectories among individuals and family groups, in interbirth intervals depending on offspring’s fate, and consequently on lifetime reproductive success for the caring parent (Clutton-Brock 1991).

Capture-recapture (CR) models allow studying species with complex demography in the wild, e.g. by considering ‘breeder’ and ‘non-breeder’ reproductive states to estimate breeding probabilities and status-specific demographic parameters while accounting for imperfect detectability (e.g., Lebreton et al. 2009). One can distinguish between successful and failed breeding events (e.g., Lagrange et al. 2017) and include varying litter or clutch size (e.g., Doligez et al. 2002) and memory effects (Cole et al. 2014), to investigate the costs of reproduction on survival and future reproduction for species providing short-term parental care, i.e. when offspring reach independence before the next breeding season (e.g., Yoccoz et al. 2002). Indeed, most CR models rely on the assumption of independence among individual CR histories (Lebreton et al. 2009).

In the case of species providing EPC, one challenge stems from the multiple-year dependency among individual’s life histories within parent-offspring units. Only few attempts have been made to tackle this issue when estimating demographic parameters, despite the fact that species providing EPC are often among long-living mammals vulnerable or threatened with extinction (e.g. polar bears, orangutans, elephants). Lunn et al. (2016) and Couet et al. (2019) proposed to model CR histories of family units (instead of individuals) which permits to consider the multiple-year dependency of offspring survival upon mother survival status and of female breeding probability upon offspring survival status. Couet et al. (2019) provided estimates of dolphin reproductive parameters corrected for state uncertainty but their model assumed a fixed age and timing at offspring independence.

Another challenge involves dealing with uncertain timing at offspring independence, when the offspring departs the caring parent(s) and becomes independent. When studying free-ranging populations, this key life history event is rarely directly observed. When a mature individual is observed without dependent offspring, it is often impossible to know if its offspring have died or already departed its natal group. As a result, estimates of demographic rates and trade-offs can be underestimated. Lunn et al. (2016)’s model for polar bears in Hudson Bay (later used in Regehr et al. (2018)) included variable age at independence but variability in the timing at offspring independence was not fully dealt with. Demographic rates were corrected by the average annual probability that independence occurred prior to sampling for all offspring, and offspring survival was assumed independent of litter size (Regehr et al. 2018). In most species, timing at offspring independence is variable and could depend on the offspring phenotypic traits (e.g. body size in brown bears, Dahle and Swenson 2003), on parental traits (e.g. parent-offspring conflict in kestrels, Vergara et al. 2010), social and mating system (e.g. helping behavior in humans, Kramer 2005), or other environmental determinants (e.g. food supply, Eldegard and Sonerud 2010). To our knowledge, no model is available to tackle both the issues of multiple-year dependency among individuals and variable timing at offspring independence. Because of these methodological challenges, the population-level consequences of EPC remain to be understood, especially in free-ranging animal populations.

Here, we develop a CR model specifically for species providing EPC. It is designed to handle multiple-years statistical dependency (until offspring independence) among individual demographic parameters by modeling CR histories grouped within family units. The model accounts for uncertain timing at offspring independence. In addition, our model allows for variability in the number of offspring born and recruited at each breeding event, variable offspring survival depending on number of siblings, and includes the influence of past reproductive history on the caring parent current status. Finally, estimates of survival rates, breeding probability and litter size probability are corrected for imperfect detection possibly depending upon family unit composition.

In what follows, we present the model, assess its performances using simulated data, and illustrate its use with a long-term dataset collected on the Svalbard polar bears. Female polar bears rely solely on stored fat reserves during pregnancy and the first three months of lactation, before feeding and protecting litters of one to three young, usually during two more years (Ramsay and Stirling 1988). They can lose more than 40% of body mass while fasting (Atkinson and Ramsay 1995). In many areas, climate change and related sea ice decline impact female bear condition and capacity to provide care for their young, with an associated decline in reproductive output (Derocher et al. 2004; Stirling and Derocher, 2012, Laidre et al. 2020). More insights into the species demography, such as the consequences of long-duration parental care on mother and offspring life histories, could help our understanding of polar bear population responses to environmental perturbations and extinction risks in future decades (Hunter et al., 2010; Regehr et al., 2016).

## METHODS

### 1. Capture-recapture model for species providing EPC

#### 1.1 Principle

We develop a CR model in the multievent framework (Pradel 2005) that is also known as a hidden Markov modeling framework (Gimenez et al. 2012). The principle is to relate the field observations, called events, to the underlying demographic states of interest through the observation process. Uncertainty on state assignment due to variable timing at offspring independence is included in the observation process. In parallel, the state process describes the transition rates between states from one year to the next. The transition rates correspond here to the demographic parameters corrected for imperfect detection and state uncertainty. Below we describe the general procedure to specify the model by defining the states and state-to-state transition process, then the events and observation process. However, for simplicity, the events and states are chosen to match the polar bear life cycle. The resulting model assumptions and its applicability to other species are discussed below.

#### 1.2 Specification of states and state process

One specificity of our model lies in the use of CR histories based on family groups instead of individuals, which permits to include the multiple-year dependency among the caring parent and dependent offspring’ demographic rates and life history traits.

States correspond to the ‘real’ demographic states of the individuals composing the family. We consider 12 states S, *S={J2,J3,SA4,SA5,A01,A02,A11,A12,AS1,AS2,A,D}*, to represent the polar bear life cycle (defined in Table 1). In addition, we specify 13 intermediary states S’, S’={J3, SA4, SA5, A11, A12,AS1,AS2,A0-,A1-,I/AS1,I/AS2,A,D}, and 16 intermediary states S’’, S’’={J3,SA4,SA5,A11,A12,AS1,AS2,B/A0-,NB/A0-,B/A1-,NB/A1-,B/AS,NB/AS,B/A,NB/A,D} (defined in Table 1). The specification of intermediary states is what permits to distinguish between failed and successful breeders in the transition matrix to consider the influence of past reproductive history on parameters (see below).

**Table 1.**
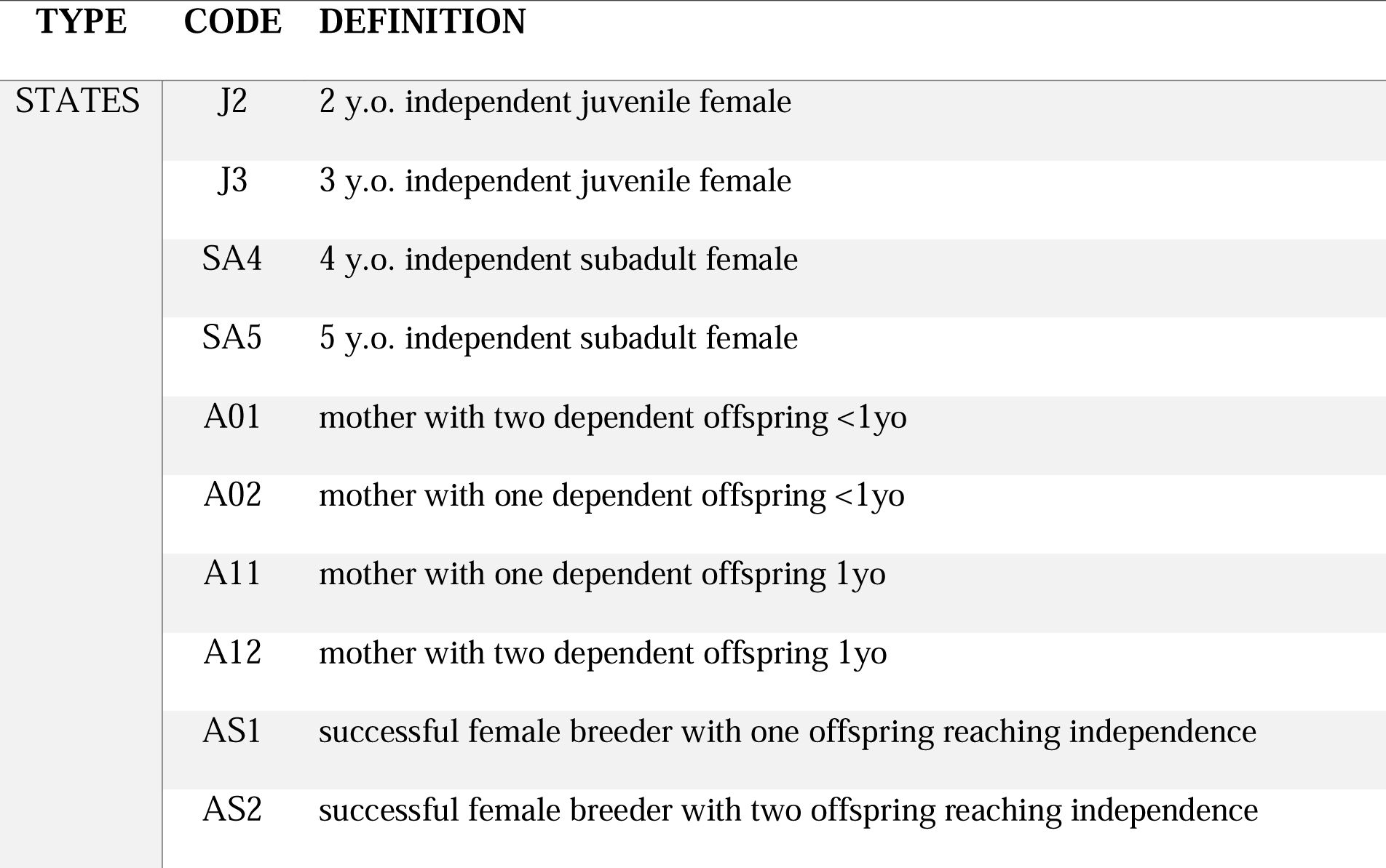
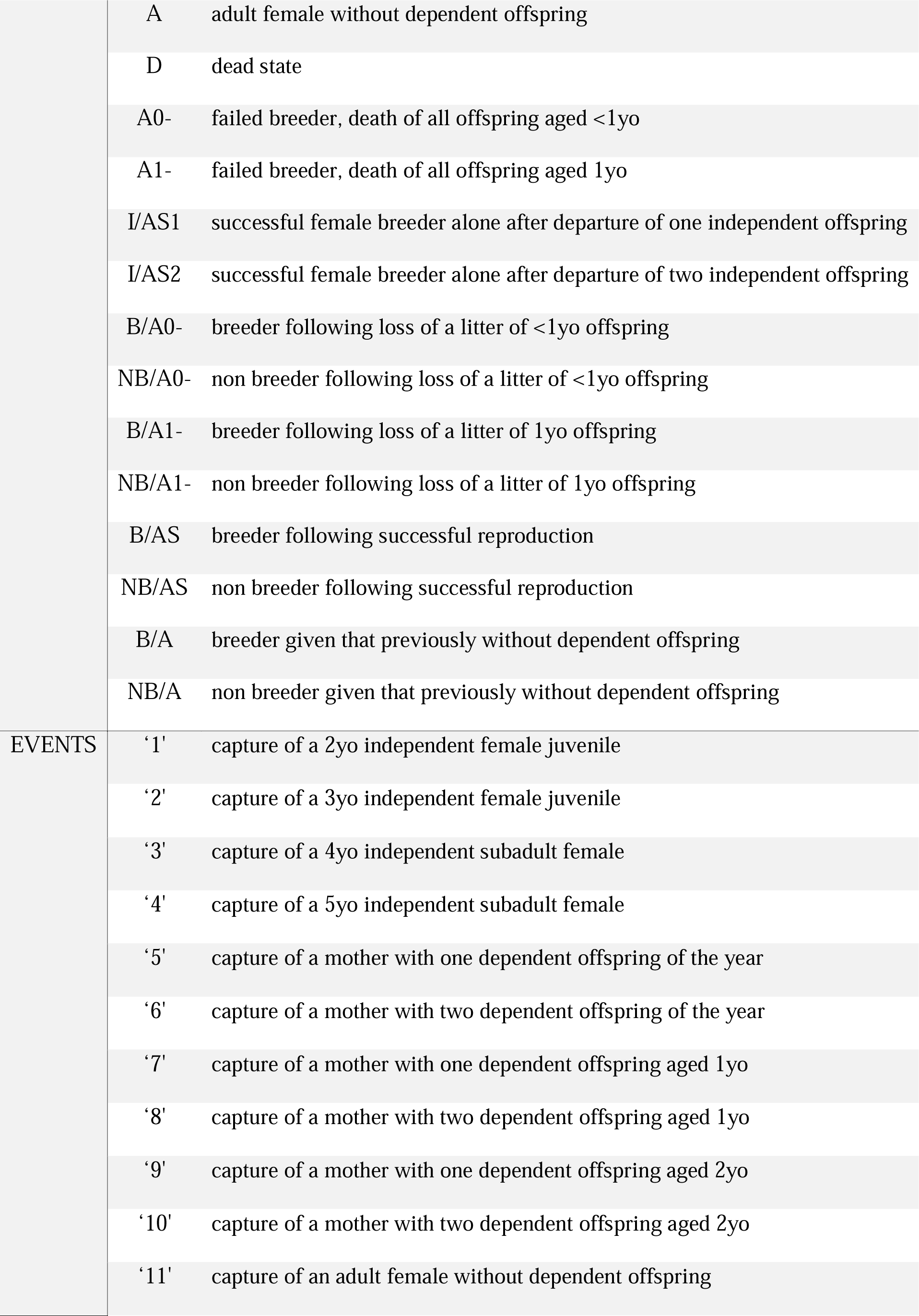
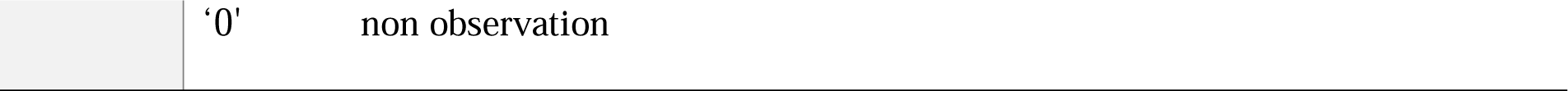
Definition of the states and events used in the model to describe the polar bear life cycle. ‘0’non observation

The model is conditioned upon first capture. The initial state vector, s_0_, gathers the proportions of family units in each state S at first capture, *s*_0_ = π_1_,…, π_11_, 0)’ (with π_12_ = 0 for state D, because an individual must be alive at first capture).

The transition matrix, Ψ, describing all possible state-to-state transitions from spring one year (t) to spring the next year (t+1), is obtained as the matrix product of four matrices Ψ = Φ · Ψ_1_ · Ψ_2_ · Ψ_3_. This decomposition is another particularity of our model which permits to estimate the relevant set of demographic parameters: independent juvenile, subadult and adult survival (matrix Φ), dependent offspring survival (matrix Ψ_1_), breeding probabilities (matrix Ψ_2_) and litter size probabilities (matrix Ψ_3_), and potential trade-offs among them. This formulation of the transition matrix implies that litter size is conditioned upon breeding decision, itself conditioned upon offspring survival, itself conditioned upon survival of the caring parent to deal with the statistical dependency existing among individuals within family units.

The Φ matrix (eq. 1) describes transitions from each state S at time t (rows) to each state S after the occurrence of the survival process for independent individuals (columns):

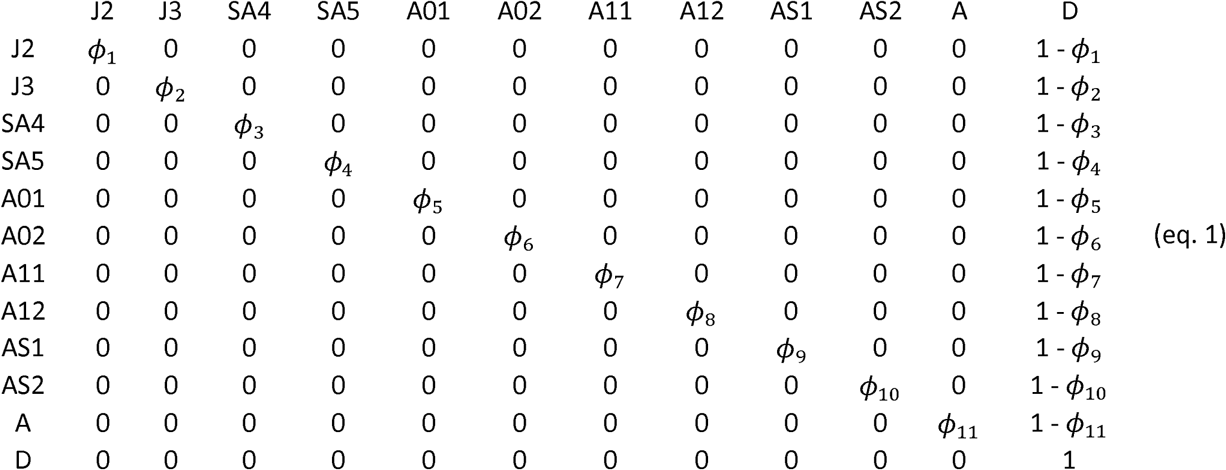

In the Φ matrix, *ϕ*_1_, …, *ϕ*_11_ correspond to survival of immature independent (juveniles and subadults) and adult female bears.

The Ψ_1_ matrix (eq.2) describes transitions from states S after the occurrence of the survival process for independent individuals (rows) to states S’ after the occurrence of the offspring survival process (columns):

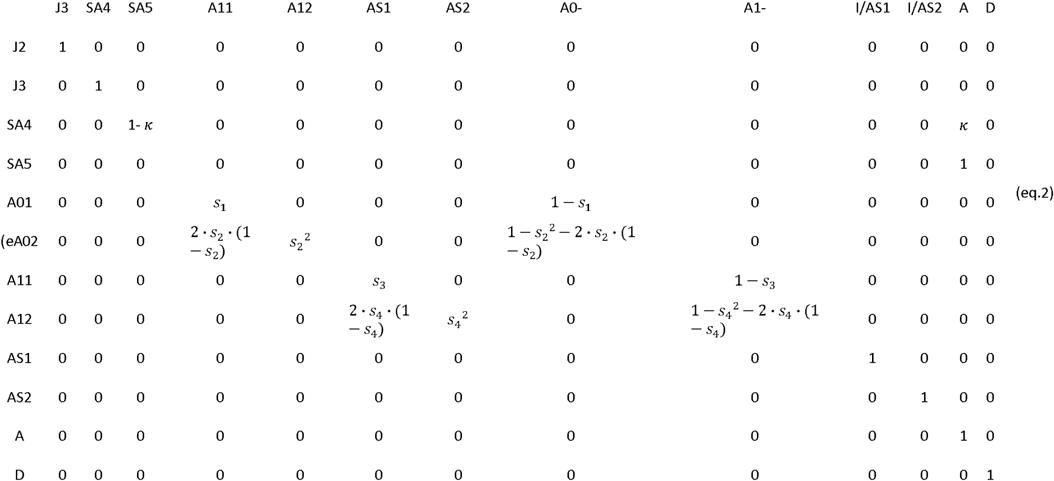

In the Ψ_1_ matrix, *κ* is the probability of first reproduction at age 5, *s* is dependent offspring survival conditioned upon mother survival (*s*_1_ and *s*_2_ for singleton cub resp. yearling; *s*_3_ and *s*_4_ for twin litter’s cub resp. yearling). Litter survival rates can be obtained from individual offspring survival rates (for singleton litters *l*_01_ = *s*_1_ and *l* _11_ = *s*_3_ for cub resp. yearling, and for twin litters *l*_02_ = 1 − (1 − *s*_2_^2^ − 2 ·*s*_2_ · (1 − *s*_2_)) and *l* _12_ = 1 − (1 − *s*_4_^2^ − 2·*s*_4_ ·(1 − *s*_4_)) for cub resp. yearling).

The Ψ_2_ matrix (eq.3) describes transitions from states S’ after occurrence of the survival processes (rows) to states S’’ depending on breeding decision (columns):

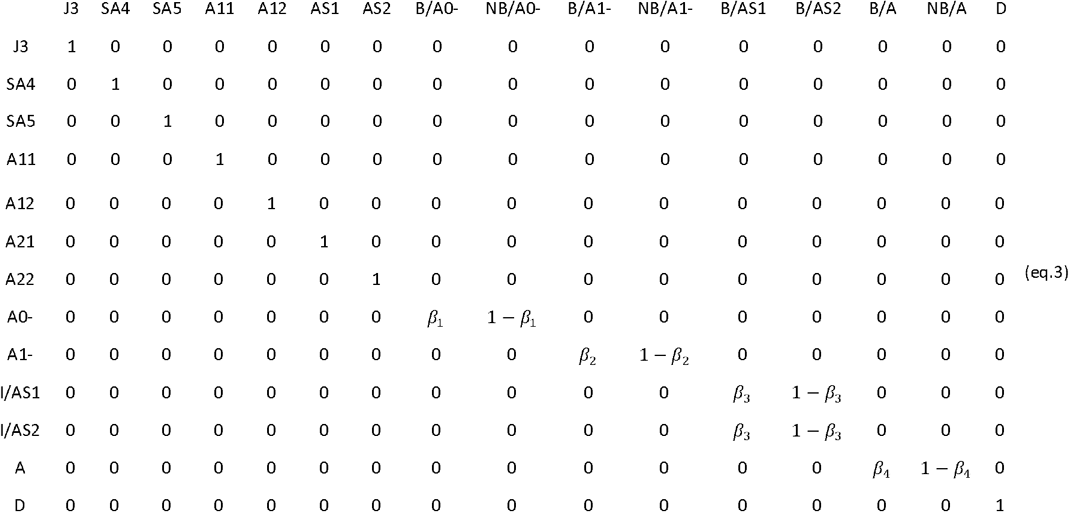

Parameter *β* is breeding probability conditioned upon mother and offspring survival status (*β*_1_ and *β*_2_ following the loss a cub litter resp. loss of a yearling litter for failed female breeders, *β*_3_ for successful breeder, *β*_4_ for female without dependent offspring at the beginning of the year).

The Ψ_3_ matrix (eq. 4) describes transitions from states S’’ after occurrence of the survival processes and breeding decision (rows) to states S at t+1 after determination of litter size for breeders (columns):

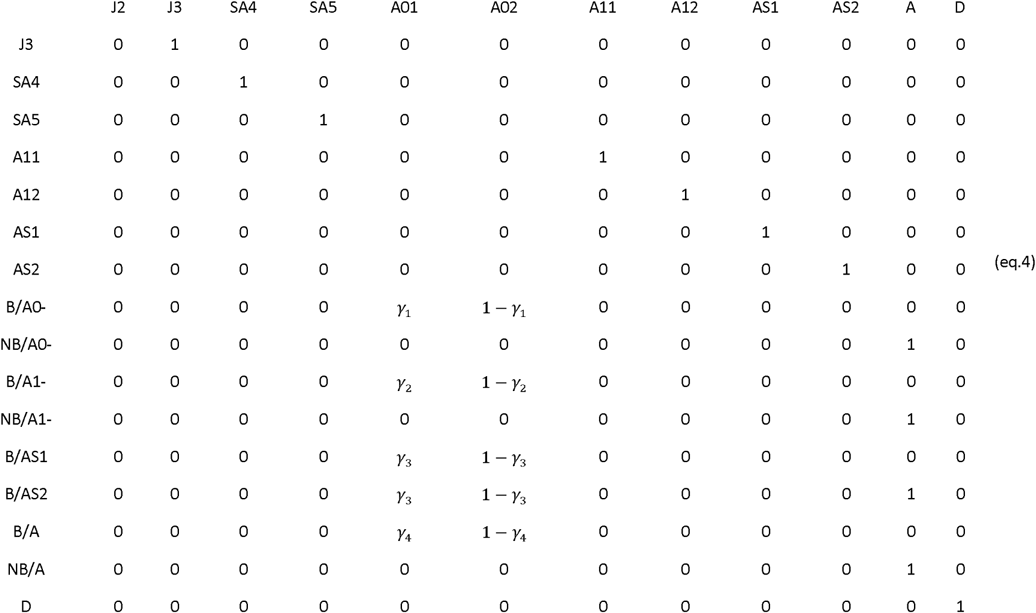

Parameter *γ* is the probability of producing a singleton litter conditioned upon mother’s and offspring’s survival status and upon breeding decision (*γ* _1_ and *γ*_2_ following the loss a cub litter resp. loss of a yearling litter, *γ*_3_ for successful breeder, *γ*_4_ for female without dependent offspring at the beginning of the year).

By modifying the constraints on parameters (i.e. setting them equal or different among states), the model can be used to investigate: i) the cost of reproduction on parent survival (by comparing the ϕ’s in matrix Φ), ii) the influence of litter size on individual offspring survival (by comparing *s*_1_ to *s*_2_, and *s*_3_ to *s*_4_ in matrix Ψ_1_) and on litter survival (by comparing l_01_ to l_02_ and l_11_ to l_12_), iv) the influence of past reproductive history on breeding probability (by comparing the *β* ‘s among them in matrix Ψ_2_), and on litter size probability (by comparing the *γ* ‘s among them in matrix Ψ_3_).

#### 1.3 Specification of events and observation process

The events correspond to the observation or non-observation of family units in the field at each sampling occasion. Each event is coded depending on the number and age of the individuals composing the family. Here, we consider 12 possible events, Ω = {1,2,3,4,5,6,7,8,9,10,11,12} to describe field observations for polar bears family groups (defined in Table 1).

In a multievent model, one specific event may relate to several possible states. Due to variable timing at offspring independence, a female successful breeder (state ‘AS1’ or ‘AS2’) can be captured together with (event ‘9’ or ‘10’) or without (event ‘11’) its two-year old offspring or not captured (event ‘12’) depending on i) whether the offspring has already departed from its mother at the time of capture and ii) on capture probability. To include uncertainty on state assignment due to variable timing at offspring independence, we decompose the observation process into two event matrices, *E*_1_ and *E*_2_, modeling respectively departure probability (*α*) and capture probability (p).

The *E*_1_ matrix (eq. 5), relates the states S to the possible observations at the time of capture, O = {1,2,3,4,5,6,7,8,9,10,11,12} (same code as the events, see Table 1), through the departure probability, denoted *α*.

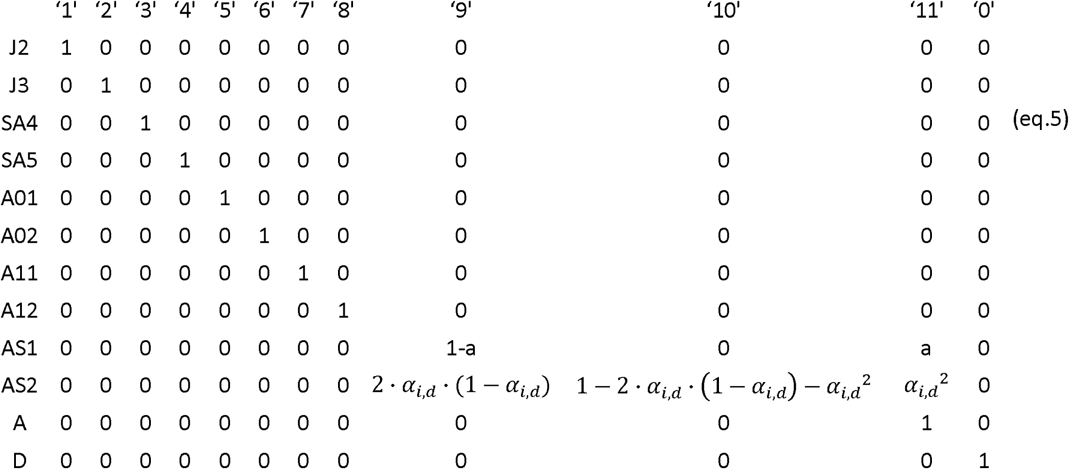

Here, *α*_*i,j*_ is the departure probability of a two-year old individual offspring belonging to family unit i on its date of capture d. The relationship between date of capture and departure probability is species-specific and either be assessed from prior knowledge on the species’ biology or field data (see Appendix 2). Here we assume that siblings’ timing at independence can, but does not have to, occur independently (if both offspring can only depart the family on the same date, the transition from state ‘AS2’ to event ‘9’ should be set to 0).

The second event matrix, *E*_2_, (eq. 6) relates all possible observations at the time of capture O to the events, Ω, actually observed in the field through the state-dependent capture probability, denoted p_s_.

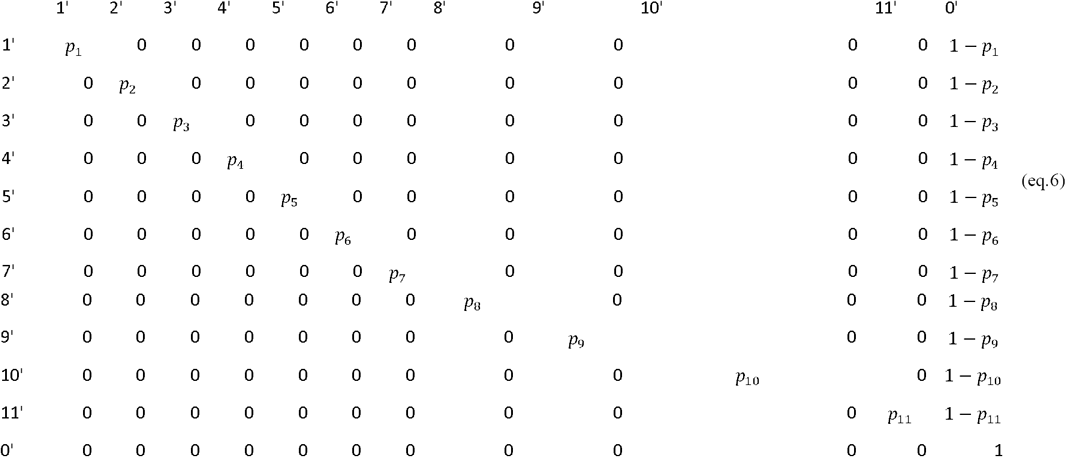

The composite event matrix, E, which relates the events to the states, is obtained as the matrix product of these two matrices *E = E*_1_ · *E*_2_. Because the model is conditioned upon first capture, the initial event vector, *e*_0_, takes the value of 1 for all events (except of 0 for the event ‘12’ corresponding to a non-observation).

#### 1.4 Applicability to other species

The model described above matches the polar bear life cycle. The model therefore assumes that care is provided by the mother only, to one or two offspring (we ignored triplets present at a low probability in polar bears), for a duration of two years maximum. A mother caring for offspring cannot mate and produce a second litter. Independent males (that have already departed from the family unit) are not included in the model.

To apply the model to other species, one should modify the number of events and states (depending on age at sexual maturity, maximum litter size, duration of parental care) and the shape of the relationship between departure probability and date of capture to match the species life cycle. In species like primates or wolves, departure from the family unit can occur throughout the year, at a more or less constant rate depending on the season, while it occurs only between February and May in polar bears due to environmental constraints. The general model structure, types of parameters involved and steps of model definition should remain similar. The influence of individual traits such as age or body weight, or environmental variables such as temperature, can be included in the model under the form of individual or temporal covariates (Pollock 2002). In addition, specificities related to data collection can also be included in a similar way, such as trap effects (Pradel and Sanz-Aguilar 2012) or latent individual heterogeneity by using mixture of distributions or random effects (Gimenez et al. 2018). Guidance to fit the model in a Bayesian framework in program Jags for real and simulated data are provided in Appendices 1 and 2.

##### Simulation study

The simulation study was aimed at evaluating the performance of our model to estimate demographic parameters under various assumptions about the timing at offspring independence (constant versus seasonal departure rate) and various degrees of capture probability (low p= 0.25, high p=0.7), for a medium-size data set (T= 15 sampling occasions, with R=80 newly marked at each occasion in equal proportion among the 11 alive states).

We simulated data for a virtual long-lived mammal species mimicking the polar bear using the model described in box 1. We used *φ=* 0.9 for independent female bear survival (aged 2+ y.o.), *s*_1_ = *l*_01_ = 0.6 for singleton cub survival, *s*_2_ = 0.55 for twin litter’ s cub survival (corresponding to twin litter survival *l*_02_ = 0.7975), *s*_3_ = 0.8 for singleton yearling survival and *s*_4_ = 0.75 for twin litter’s yearling survival (corresponding to litter survival *l* _12_ = 0.94). Offspring survival rates were conditioned upon mother survival. If a mother dies, its dependent offspring had no chances of surviving. For breeding probabilities, we used *β* _1_ = 0.5, *β*_2_ = 0.7, *β*_3_ = 0.9 and *β*_4_ = 0.8. For litter size probabilities, we used *γ*_1_ = 0.4, *γ*_2_ = 0.5, *γγ*_3_ = 0.6 and *γ*_4_ = 0.7. We set *κ* = 0 and assumed that females had their first litter at age 6 or older.

We assumed that captures occurred each year between mid-March to end of May (day of the year d=80 to d=130). For each capture event, date of capture was randomly sampled from the distribution of the polar bear data dates of capture (see Appendix 1). In the constant scenario, we assumed that two-year old bears reached independence at a constant departure rate (*α*) during the field season, independently of the date of capture. We chose an intermediate value of *α* = 0.5 (if independence occurred always after the field season, *α* = 0, versus always before the field season, *α* = 1). In the seasonal scenario, we assumed that departure rate varied with date of capture (d) following a logistic relationship (regression coefficients were estimated from the polar bear data, see Appendix 2). Most of the two-year old offspring were captured with their mother at the beginning of the field season, while departure probability increased logistically up to 80% at the end of the field season.

We simulated 100 CR datasets for each of the 4 scenarios (S1: low detection with constant departure, S2: low detection with seasonal departure, S3: high detection with constant departure, S4: high detection with seasonal departure). We simulated the data using program R. We fitted the model using program jags called from R (Plummer 2016). Appendix 1 containing guidance, R code and files to simulate data and fit the model is available on GitHub at https://github.com/SCubaynes/Appendix1_extendedparentalcare.

##### Case study: Polar bears in Svalbard

In polar bears, care of offspring is provided by the mother only (Amstrup, 2003). Males were therefore discarded from our analysis. Adult female polar bears mate in spring (February to May, Amstrup 2003), and in Svalbard usually have their first litter at the age of six years, but some females can have their first litter at five years (Derocher 2013). They have delayed implantation where the egg attaches to the uterus in autumn (Ramsay and Stirling 1988). A litter with small cubs (ca 600 grams) is born around November to January, in a snow den that the mothers dig out in autumn, and where the family stay 4-5 months. The family usually emerges from the den in March-April, and stay close to the den while the cubs get accustomed to the new environment outside their home, for a few days up to 2-3 weeks (Hansson and Thomassen 1983). Litter size in early spring vary from one to three, with two cubs being most common, three cubs in most areas being rare, and commonly around one out of three litters having one cub only (Amstrup 2003). In Svalbard, polar bears become independent from their mother shortly after their second birthday (average age at independence is 2.3). Two-year old bears typically depart from the mother in spring (between mid-March to end of May), when the mother can mate again. There is only one anecdotic record of a yearling alive without his mother. Because the field season can last for several weeks, some two-year-old bears were captured together with their mother and others were already independent at the time of capture. The minimum reproductive interval for successful Barents Sea polar bears is 3 years. On the contrary, loss of a cub litter shortly after den emergence may mean the mother can produce new cubs in winter the same year (Ramsay and Stirling 1988).

Bears captured in Svalbard are shown to be a mixture of resident and pelagic bears (Mauritzen et al. 2002). To focus on the resident population, independent bears captured only once were not included in our analysis. We therefore analyzed N = 158 encounter histories of resident polar bear family units captured each spring after den emergence between doy 80 to 130 (mid-March to mid-May), from 1992 to 2019, in Svalbard. It corresponds to 81 capture events of juvenile and subadults, 231 cubs, 96 yearling, 23 dependent two-year old, and 444 captures of adult females. Polar bears were caught and individually marked as part of a long-term monitoring program on the ecology of polar bears in the Barents Sea region (Derocher 2005). All bears one year or older were immobilized by remote injection of a dart (Palmer Cap-Chur Equipment, Douglasville, GA, USA) with the drug Zoletil® (Virbac, Carros, France) (Stirling et al. 1989). The dart was fired from a small helicopter (Eurocopter 350 B2 or B3), usually from a distance of about 4 to 10 meters. Cubs of the year were immobilized by injection with a syringe. Cubs and yearlings were highly dependent on their mother; therefore, they remained in her vicinity and were captured together with their mother. A female captured alone was considered to have no dependent offspring alive. Death of the cubs could have occurred in the den or shortly after den emergence but before capture. Hereafter, estimated cub survival thus refers to survival after capture. Infant mortality occurring before capture will be assigned to a reduced litter size. Because only 3% of females were observed with 3 offspring, we analyzed jointly litters of twins with triplets.

We built the model described above with 12 states and 12 events to describe the life cycle (Table 1). Preliminary analyses suggested that mother survival did not vary according to state, we therefore constrained parameter cp to be equal among all states in matrix Φ. To avoid identifiability issues due to a relatively small sample size, we assumed breeding probability and litter size probability did not vary between successful breeders (states AS1 and AS2) and female without dependent offspring (state A) by setting *β*_3_ = *β*_4_ and *γ*_3_ = *γ*_4_.. We also assumed that litter size probability did not vary among failed breeders (loss of cub versus yearling litter), by setting *γ*_1_ = *γ*_2_. We could not assess formally the fit of our model because no test is yet available for multievent models.

Using the conditional probabilities estimated in the model, we calculated the net probability for a female to raise none, Pr(X=0), one, Pr(X=1), or two offspring, Pr(X=2) to independence over a 3-year period (details are provided in Appendix 2).

We considered adult females without dependent offspring at the beginning of the time period, so that we have:

Appendix 2 containing guidance, R code, data files to fit the model to the polar bear data, and additional results is available on GitHub at https://github.com/SCubaynes/Appendix2_extendedparentalcare.

## RESULTS

### Model performance evaluated on simulated datasets

Model performance was satisfying and comparable in all 4 simulated scenarios (S1 with low detection and constant departure, S2 with low detection and seasonal departure, S3 with high detection and constant departure and S4 with high detection and seasonal departure), with low average bias (*B*_*S*1_ = - 0.000, *B*_*S*2_ = - 0.004, *B*_*S*3_ = - 0.004, *B*_*S*4_ = - 0.003) and root-mean-square error (*rmse*_*S*1_ = 0.042, *rmse*_*S*2_ = 0.041, *rmse*_*S*3_ = 0.031 and *rmse*_*S*4_ = 0.031) (see Appendix 1 for details).

For most parameters, bias was very low, B < 0.02, except for parameters β_2_ in scenarios S2, S3 and S4 (−0.04 < B < -0.03) and *s*_3_ in scenario S2 (B = -0.03), rmse < 0.05, except for parameters *β* _2_, *γ* _1_ and *γ* _2_ (0.05 < rmse < 0.07). For these three parameters, precision was lower in the two scenarios with low detection (see Appendix 1). Estimates obtained for the scenario mimicking the polar bear study case (S2: low detection and seasonal departure) are provided in Figure 2.

**Figure 1:**
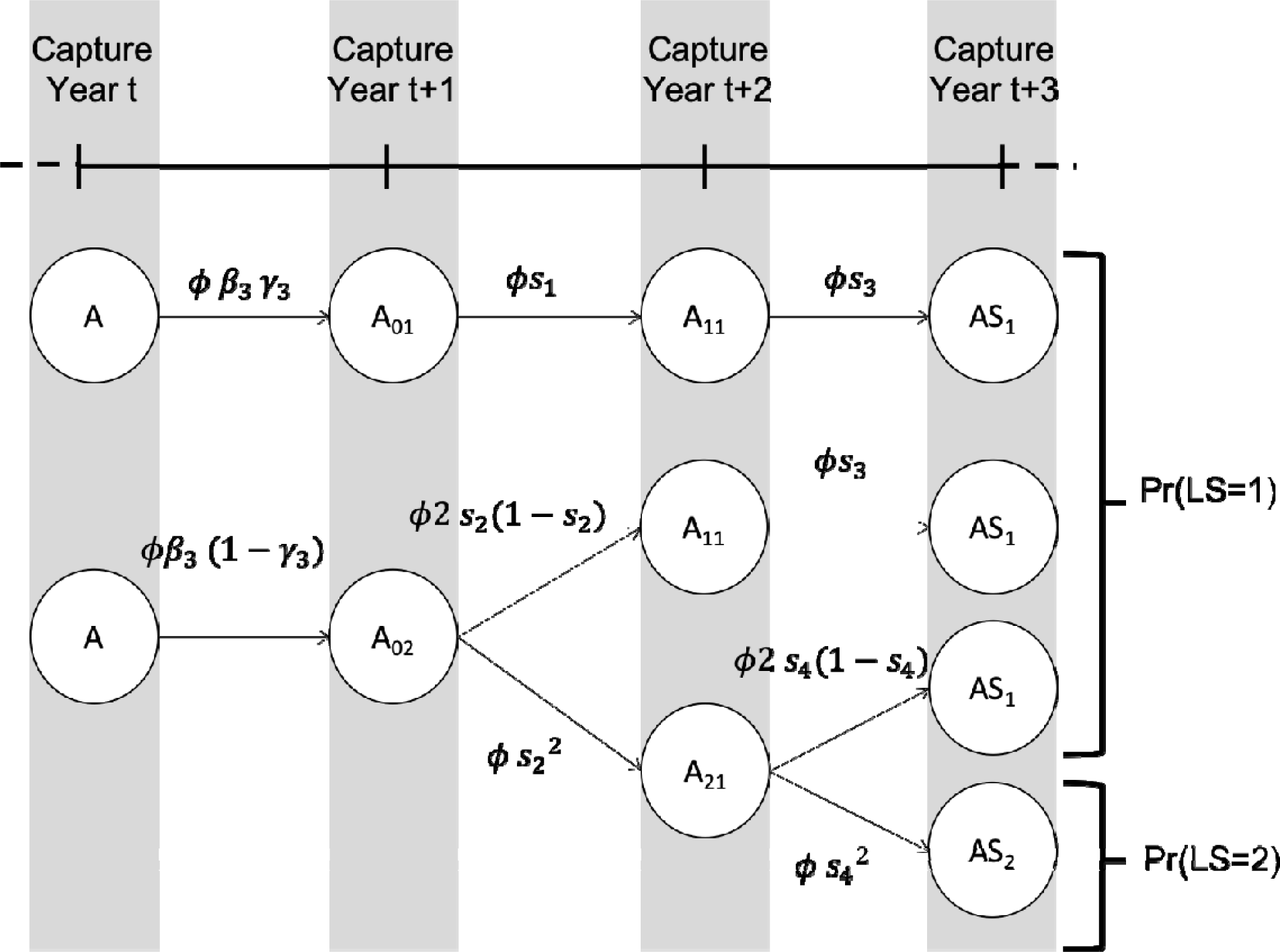
Life history events with associated probabilities of raising one (X=1) or two (X=2) offspring to independence over a 3 years period for a female polar bear alive and without dependent offspring at the beginning of the period (is adult survival, is breeding probability of a female without dependent offspring, is the probability of a singleton litter, and resp. cub and yearling survival in a singleton litter, and resp. cub and yearling survival in a twin litter).

**Figure 2.**
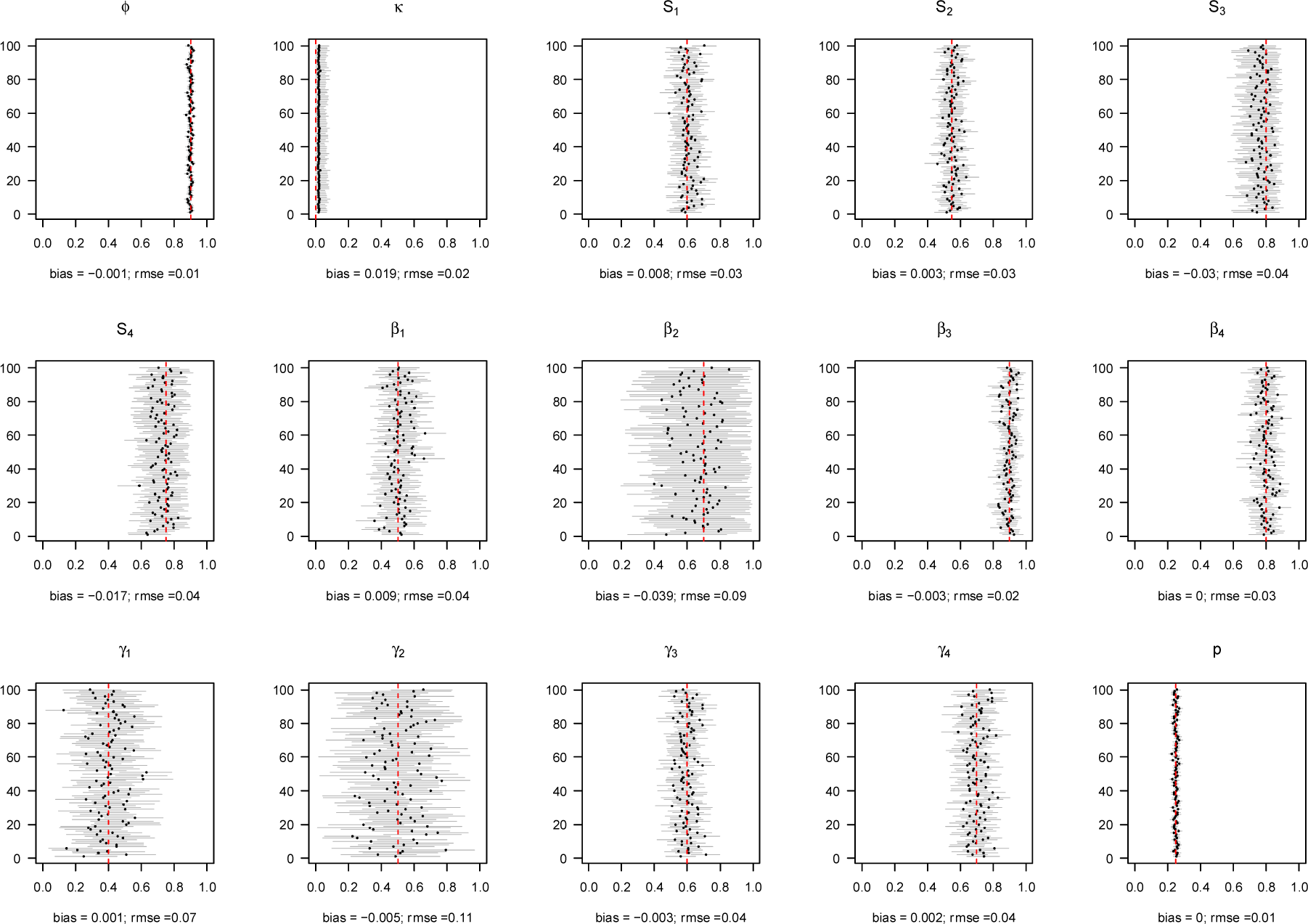
Performance of the model on simulated data with low detection with seasonal departure (scenario S2). For each of the 100 simulated data sets, we displayed the mean (circle) and the 95% confidence interval (horizontal solid line) of the parameter. The actual value of the parameter is given by the vertical dashed red line. The estimated absolute bias and root-mean-square error are provided in the legend of the X-axis for each parameter. Regarding notations, *ϕ* stands for juvenile, subadult and adult survival, *κ* is the probability of first reproduction at age 5, *s* is dependent offspring survival conditioned upon mother survival (*s*_1_ and *s*_2_ for singleton cub resp. yearling; *s*_3_ and *s*_4_ for twin litter’s cub resp. yearling), *β* is breeding probability conditioned upon mother and offspring survival status (*β* _1_ and *β*_2_ after loss a cub litter resp. loss of a yearling litter, *β*_3_ for successful breeder, *β*_4_ for female without dependent offspring), *γ* is the probability of producing a singleton litter conditioned upon mother’s and offspring’s survival status and upon breeding decision (*γ*_1_ and *γ*_2_ after loss a cub litter resp. loss of a yearling litter, *γ*_3_ for successful breeder, *γ*_4_ for female without dependent offspring), *p* is detection probability.

### Case study: Polar bear demography

Departure probability was about 40% at the end of March and reached 80% at mid-May (Appendix 2). About half of the two-year old bears had departed their mother at the time of capture. Estimates of demographic parameters are provided in Table 2 (more results are provided in Appendix 2). Independent female (aged 2+) survival was high (0.93). Individual offspring survival rates, conditioned upon mother survival, did not vary significantly with litter size for cubs or yearlings. Average yearling survival was lower for singleton (0.67) than for litters of twin (0.80), although the 95% credible intervals did overlap. Concerning litter survival conditioned upon mother survival, it was higher for twin compared to singleton, for both cubs’ and yearlings’ litters. A small proportion of females, about 12%, started to reproduce (i.e. produced a litter that survived at least until the first spring) at 5 y.o. Outcome of the previous reproduction influenced breeding probability. Breeding probability following the loss of a cub litter during the year (after capture) was low, about 10%, while it was about 50-60% for female’ successful breeders or without dependent offspring at the beginning of the year or after the loss of a yearling litter. Detection probability was relatively low, about 0.25 (0.22 - 0.27). At first capture, 37% were independent juvenile or subadult females, 18% were adult females alone, 28% were adult females with one or two cubs, 12% with one or two yearlings and 5% with two-year old bears.

**Table 2:**
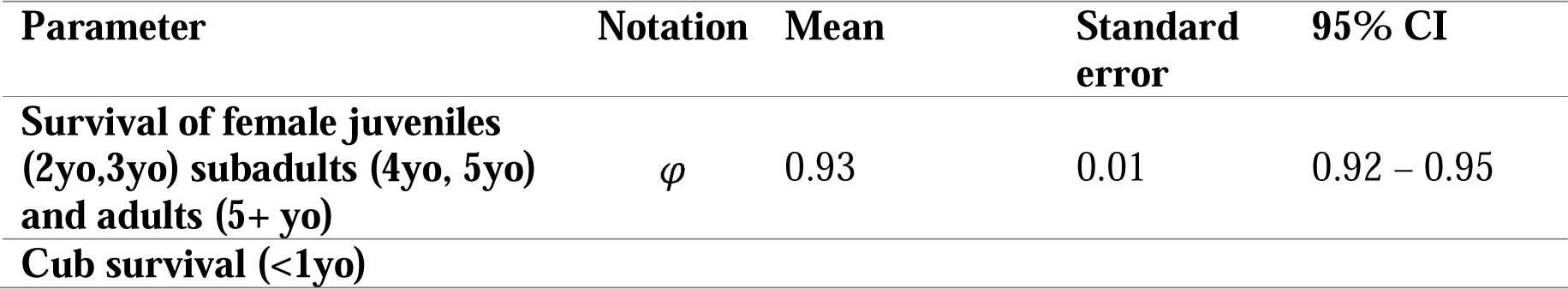
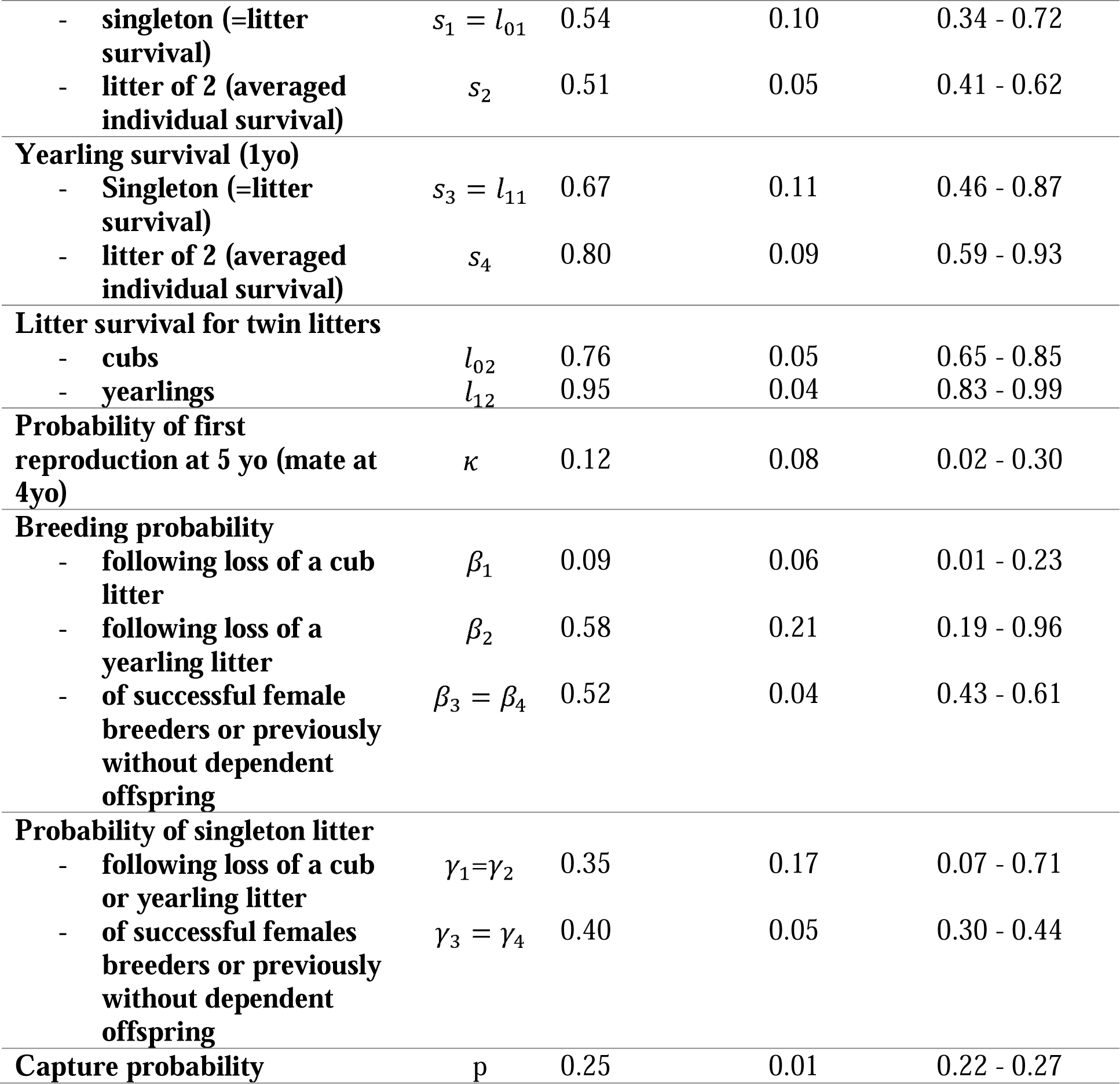
Parameter estimates. Means are given with 95% credible intervals (CI). Dependent offspring (cub age <1 y.o., yearling 1 y.o.) survival and breeding probabilities are conditioned upon mother survival, litter size probability of producing a singleton is as well conditioned upon breeding decision.

Over a 3 years period, the probability to successfully raise one offspring to independence for a female polar bear, alive and without dependent offspring at the beginning of the period, was on average 0.29 (0.20-0.38) and 0.04 (0.02-0.07) to raise two offspring to independence. The probability of failed breeding (no offspring successfully reached independence) over this period was high, about 0.67 (0.57-0.76). Note that this calculation includes breeding probability, and therefore does not reflect offspring survival until independence (see Method section).

## DISCUSSION

Overall, our model performed well in estimating all demographic parameters in all the simulated scenarios. A multievent approach is a promising tool to deal with uncertainty on the timing at offspring independence both when departure probability was constant or varied within the field season. Estimates obtained for adult and offspring survival, probability of sexual maturation, breeding probability (except after the loss of a yearling litter), litter size probabilities and detection probability were unbiased in most simulated scenarios. Precision was satisfying in most cases, but it was lower for breeding probability after the loss of a yearling litter and for litter size probabilities of failed breeders, especially in scenarios with low detection (Appendix 1). These specific parameters should therefore be interpreted with caution for the study case. In our simulations, T=15 sampling occasions appeared sufficient to obtain satisfying estimates for most parameters. In the polar bear data, there were few recaptures of females on subsequent years due to relatively low detection rate. As a result, preliminary analyses suggested a potential confusion between these parameters. We dealt with this issue by including a biologically realistic constraint on prior distributions, stating that cub survival was lower than that of yearling survival (Amstrup and Durner, 1995) which was enough to ensure parameter estimability. Inference in a Bayesian framework is useful in this regard, because it allows to include prior information when available (McCarthy and Masters 2005) to improve the estimation of model parameters.

For polar bears, we showed that outcome of the previous breeding event influenced breeding probability. Reduced offspring survival one year, for example due to poor environmental conditions (Derocher et al. 2004), might therefore increase intervals between successful reproduction through reduced breeding probability the next year (Wiig 1998). This means that by ignoring multiple-year dependency among mother and offspring, classical models can underestimate reproductive intervals, therefore risking to overestimate the population growth rate. However, the biological relevance of our model is currently limited, because we ignored temporal and individual heterogeneity among females in the model. Survival rates for independent female bears (0.93) were close to an earlier study for the same population (0.96), based on telemetry data (Wiig 1998). Our results may overestimate dependent offspring survival because we focused on resident bears captured more than once. Wiig (1998) results indicated that females in Svalbard went into den on average every second year (while successful breeding means denning on a three-year interval), which seems coherent with our results (about 33% chances of successful reproduction over a three-year period).

Here, we proposed a general model structure that can be applied to other species providing EPC. The originalities of our approach lie in using family structure to define statistical units in our model, and the inclusion of variable timing at offspring independence. It allows to include dependency among individuals over multiple-years and therefore evaluate correlations between offspring and parents’ life history parameters. Our model could be applied to other species providing EPC, such as wolves, showing variable litter size and age at offspring independence. However, applying our model to species like elephants, killer whales or humans would require including an additional ‘post-reproductive’ stage and estimate the age at menopause. Such extensions of our model could be used, for example, to evaluate the population-level consequences of positive or negative correlation between parents and offspring traits (e.g. food sharing among group members Lee 2008; or parent-offspring conflict Kölliker et al. 2013). For polar bears specifically, future analyses will integrate in the model the effect of female age on survival and reproductive success (Atkinson and Ramsay 1995; Folio et al. 2019) and influence of climatic variables on body weight and demography (Derocher et al. 2004; Stirling and Derocher 2012) as individual and environmental covariates in a regression-like framework. Our model could then be used to provide population predictions of the demographic response of the Barents Sea polar bear population under climate change (Hunter et al., 2010; Regehr et al., 2016, 2018, Laidre et al. 2020).

## Authors’ contributions

SC, OG, NY and RP conceived the ideas and designed methodology; JA, RAI and ØW collected the data; SC analysed the data; SC led the writing of the manuscript. All authors contributed critically to the drafts and gave final approval for publication.

